# Plant cell wall integrity maintenance and pattern-triggered immunity modulate jointly plant stress responses in Arabidopsis thaliana

**DOI:** 10.1101/130013

**Authors:** Timo Engelsdorf, Nora Gigli-Bisceglia, Manikandan Veerabagu, Joseph F. McKenna, Frauke Augstein, Dieuwertje van der Does, Cyril Zipfel, Thorsten Hamann

## Abstract

Plant cells are surrounded by walls, which must often meet opposing functional requirements during plant growth and defense. The cells meet them by modifying wall structure and composition in a tightly controlled and adaptive manner. The modifications seem to be mediated by a dedicated cell wall integrity (CWI) maintenance mechanism. Currently the mode of action of the mechanism is not understood and it is unclear how its activity is coordinated with established plant defense signaling. We investigated responses to induced cell wall damage (CWD) impairing CWI and the underlying mechanism in *Arabidopsis thaliana.* Interestingly inhibitor- and enzyme-derived CWD induced similar, turgor-sensitive stress responses. Genetic analysis showed that the receptor-like kinase (RLK) FEI2 and the mechano-sensitive, plasma membrane-localized Ca^2+^- channel MCA1 function downstream of the THE1 RLK in CWD perception. Phenotypic clustering with 27 genotypes identified a core group of RLKs and ion channels, required for activation of CWD responses. By contrast, the responses were repressed by pattern-triggered immune (PTI) signaling components including PEPR1 and 2, the receptors for the immune signaling peptide *At*Pep1. Interestingly *At*Pep1 application repressed CWD-induced phytohormone accumulation in a *PEPR1/2*-dependent manner. These results suggest that PTI suppresses CWD-induced defense responses through elicitor peptide-mediated signaling during defense response activation. If PTI is impaired, the suppression of CWD-induced responses is alleviated, thus compensating for defective PTI.

**Significance statement:** Stress resistance and plant growth determine food crop yield and efficiency of bioenergy production from ligno-cellulosic biomass. Plant cell walls are essential elements of the biological processes, therefore functional integrity of the cell walls must be maintained throughout. Here we investigate the plant cell wall integrity maintenance mechanism. We characterize its mode of action, identify essential signaling components and show that the *At*Pep1 signaling peptide apparently coordinates pattern triggered immunity (PTI) and cell wall integrity maintenance in plants. These results suggest how PTI and cell wall modification coordinately regulate biotic stress responses with plants possibly compensating for PTI impairment through enhanced activation of stress responses regulated by the CWI maintenance mechanism.

## Main

Plants are frequently cited for their ability to adapt successfully to diverse environments by modifying themselves. A corner stone enabling this adaptability are the cell walls surrounding all plant cells. The walls are also essential elements of growth, development and resistance against biotic and abiotic stress, which influence food crop yield (1, 2). This is illustrated by mutations affecting cell wall signaling and metabolism, which form traits that have been used to improve the performance of staple crops like maize and rice (3, 4). The dynamic changes in cell wall metabolism responsible for adaptation have been summarily described as cell wall plasticity and been also identified as major obstacle in efforts aimed at modifying lignocellulosic biomass in a knowledge-based manner (5). The available evidence suggests that the plant cell wall integrity (CWI) maintenance mechanism forms an integral element of cell wall plasticity (6–9). This mechanism constantly monitors the state of plant cell walls and initiates adaptive changes in cell wall and cellular metabolism in response to cell wall damage (CWD) (10, 11). CWD is used here to summarily describe changes to cell wall structure or composition, which impair CWI. Such impairment can occur during pathogen infection when (pathogen-derived) enzymes break down the walls. This releases initially cell wall-derived fragments, leading to cell wall weakening / deformation / displacement relative to the plasmamembrane and can result eventually in cell bursting (due to the high turgor pressure levels prevalent in plant cells) (12, 13). The potential importance and the mode of action how cell wall fragments can activate plant defenses is exemplified by cellobiose or oligogalacturonides, which can activate plant immune responses (OGAs, fragments of pectic polysaccharides) (14, 15). Alternatively compounds inhibiting cell wall biosynthetic processes can also cause CWD by preventing production of load bearing elements, resulting in wall deformation / displacement of walls vs. plasmamembrane, possibly releasing fragments from the wall and eventually leading to cell bursting (16). These examples illustrate different modes of action for generation of CWD, highlight similarities and differences between them but more importantly indicate that both ligand- and/or mechano-perception-based CWD detection mechanisms are conceivable.

While a large number of candidate genes has been implicated in CWD perception in recent years, experimental evidence supporting their involvement exists only for a small number of them (6, 8, 17). Amongst them are the plasmamembrane-localized receptor-like kinases (RLK) THESEUS1 (THE1), MDIS1-INTERACTING RECEPTOR LIKE KINASE2 (MIK2), WALL ASSOCIATED KINASE2 (WAK2) and a putatively stretch-activated, mechano-sensitive Ca^2+^-channel (MID1-COMPLEMENTING ACTIVITY1, MCA1) (18–22). MCA1 was originally identified through its ability to partially complement a *Saccharomyces cerevisiae* strain deficient in MID1-CCH1, which is required for CWI maintenance in yeast (18, 23). Homologs of THE1 and MCA1 have been identified in *Oryza sativa* (*OsMCA1*), *Zea mays* (*NOD*) and *Marchantia polymorpha* (*MpTHE*) suggesting that the mechanism is conserved across the plant kingdom (24–26). While MIK2 belongs to subfamily XIIb of the LRR-RLKs, THE1 and WAK2 RLKs are containing extracellular domains potentially capable of binding cell wall derived ligands or cell wall epitopes (7, 8). However the binding capability has only been confirmed experimentally for WAK2 (27). Interestingly it was shown recently that THE1 and MIK2 are required for resistance to *Fusarium oxysporum* f. sp. *conglutinans* implicating CWI signaling in biotic stress responses (22). Previous work showed that CWD (caused by impairing cellulose production) causes production of callose, lignin, jasmonic acid (JA), salicylic acid (SA), reactive oxygen species (ROS), ethylene and implicated Ca^2+^-based signaling processes in CWI maintenance (11, 20, 28–31). Apoplastic ROS generation induced by CWD in *Arabidopsis* is apparently mediated by RBOHD (20). Interestingly the activity of RBOHD is regulated both by calcium-based as well as BRI1-ASSOCIATED KINASE1 (BAK1; via BOTRYTIS INDUCED KINASE1, BIK1)- mediated signaling during pattern triggered immunity (PTI) (32, 33). BAK1 acts as a co-receptor required for perception of PAMPs (<u>P</u>athogen-<u>A</u>ssociated <u>M</u>olecular <u>P</u>atterns) like flagellin or EF-Tu and DAMPs (<u>D</u>amage-<u>A</u>ssociated <u>M</u>olecular <u>P</u>atterns) like *At*Pep1. *At*Pep1 has been described previously as an enhancer of responses during pathogen infection regulated by pattern-triggered immunity (PTI) (34). It has been proposed that the CWI maintenance mechanism and PTI may constitute complementary elements of the mechanism regulating plant defense responses (35–37). However, to date only very limited experimental evidence for this concept is available and possible pathway interactions have not been investigated.

Here we investigate the responses to different types of CWD to understand the cellular events underlying CWD perception. We characterize 27 genotypes to establish the functions of candidate genes in CWI maintenance; perform a genetic analysis to assess if key CWI signaling elements belong to one or more signaling cascades and show that the signaling peptide *At*Pep1 can repress CWD-induced phytohormone production.

## Results and discussion

We used an *A. thaliana* seedling-based model system to determine how plants respond to different types of CWD and investigate further the role of turgor pressure in CWD perception (11). CWD was generated either using isoxaben (ISX), an inhibitor of cellulose biosynthesis in primary cell walls or driselase, an enzyme mix from *Basidomycetes sp.* containing cell wall degrading enzymes (cellulases, laminarinases and xylanases) (12, 16). Driselase was chosen as tool to cause CWD because the enzyme mix is similar to the enzyme cocktail released by pathogens during infection and the enzymatic digest generates cell wall fragments regardless of cell type or differentiation stage. Furthermore, high turgor pressure levels are not necessary to cause CWD by driselase and the end-result of the treatment is bursting cells (unless they are maintained in a medium containing osmoticum as in the case of protoplast generation). ISX was chosen because it only affects a certain cell type / differentiation stage, and does not directly damage the cell walls, instead it weakens them indirectly by inhibiting a biosynthetic process, making CWD occurrence dependent on high turgor levels (ie. it can be manipulated by using osmoticum like sorbitol, mannitol, etc.). Furthermore the availability of the *isoxaben resistant1-1* (*ixr1-1*) mutation (a point mutation in *CELLULOSE SYNTHASEA3*) represents an elegant experimental control (38). In addition to Col-0 and *ixr1-1* seedlings *bri1-associated receptor kinase1-5* (*bak1-5)* seedlings were also analyzed (39). The *bak1-5* allele used here is only impaired in immune response but not in brassinosteroid-dependent signaling, making it suitable for detecting effects caused either by DAMPs (e.g. *At*Pep1) generated by CWD or PAMPs possibly contaminating driselase (39). JA, SA, callose and lignin production were used as readouts since they represent established CWD and immune responses (11). The role of turgor pressure was assessed through co-treatments with an osmoticum (11). JA and SA levels were low in mock (DMSO)-treated Col-0 and *bak1-5* seedlings and slightly elevated in *ixr1-1* (Fig. 1A, B). ISX-treatment induced JA production in *bak1-5* seedlings more than in Col-0, while no induction was observed in *ixr1-1*. SA levels were similarly elevated in ISX-treated Col-0 and *bak1-5* seedlings whereas no accumulation was observed in *ixr1-1*. Co-treatments with sorbitol repressed the ISX-induced JA/SA production in Col-0 and *bak1-5*. Interestingly phytohormone levels were lower in mock/sorbitol and ISX/sorbitol treated *ixr1-1* seedlings than in mock-treated ones, suggesting that hyperosmotic treatments reduced indeed stress in *ixr1-1* seedlings. While mock-treated Col-0 and *bak1-5* seedlings did not show callose deposition in cotyledons, *ixr1-1* seedlings showed slightly elevated callose deposition (Figure 1C). ISX-treatment induced callose deposition strongly in Col-0, but significantly less in *bak1-5* seedlings with induction being in both cases sensitive to sorbitol co-treatments. In *ixr1-1* seedlings ISX-treatment did not induce callose deposition but the sorbitol treatment reduced the deposition compared to mock treatments. Driselase treatments induced JA/SA production in Col-0, *bak1-5* and *ixr1-1* seedlings compared to treatment with boiled driselase to different degrees, showing that active enzyme was required for induction (Figure 1D, E). While induction of both JA and SA by driselase was less pronounced than in ISX-treated seedlings (Figure 1A, B, D, E), sorbitol co-treatments still reduced or prevented accumulation in all genotypes examined. Callose induction was less pronounced in driselase-treated Col-0 seedlings than in ISX-treated ones but again sensitive to sorbitol co-treatment (Figure 1F). Interestingly callose deposition was not induced in driselase-treated *bak1-5* and *ixr1-1* cotyledons. A similar effect has been shown for *bak1-5* in flg22 treated plants, whereas the combination of elevated basal callose levels and limited effects of driselase treatment on callose production cause the absence of significant increases in *ixr1-1* cotyledons (40).

**Fig. 1.**
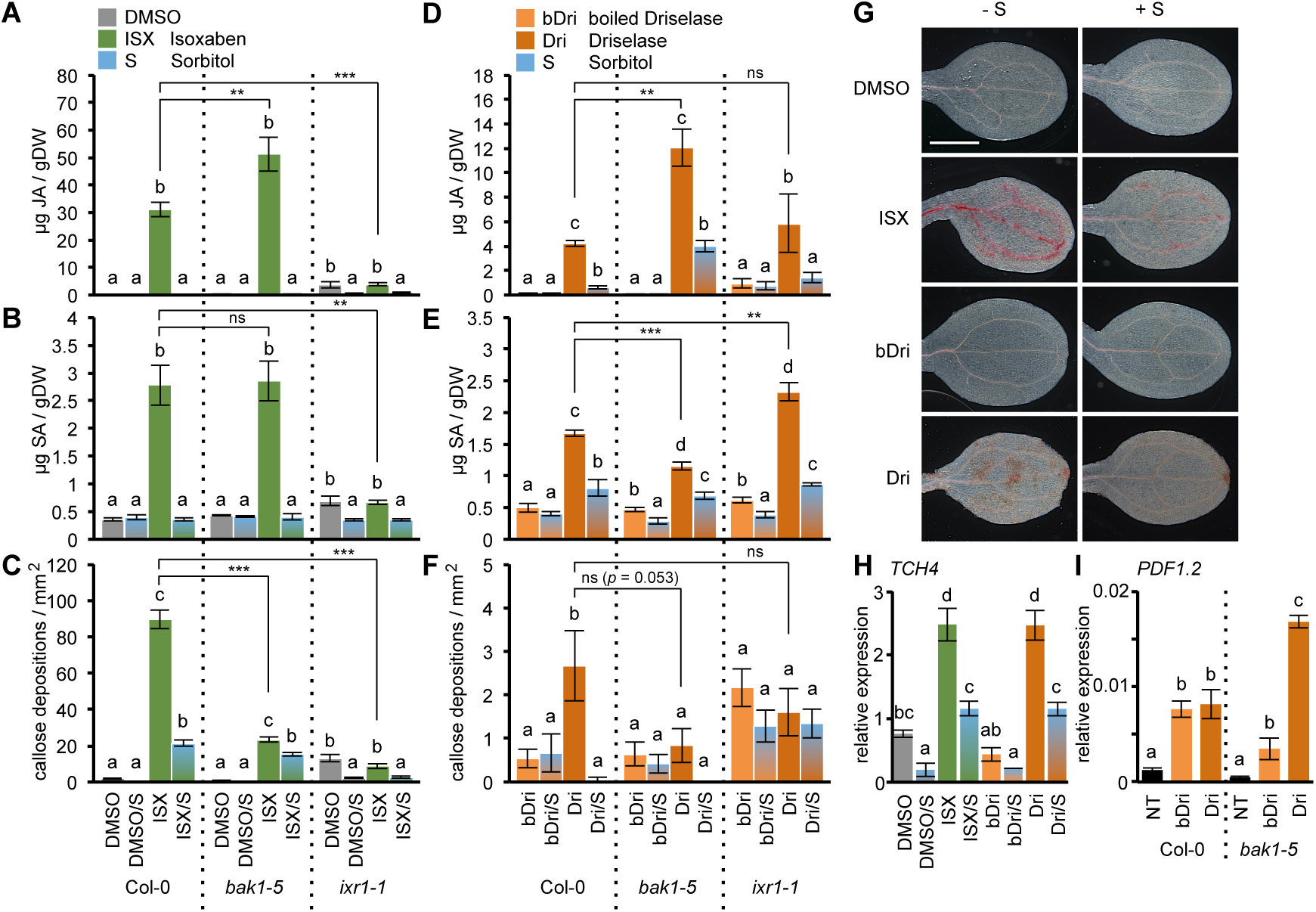
Different types of cell wall damage induce similar osmo-sensitive responses. 6 d-old Col-0, *bak1-5* and *ixr1-1* seedlings were treated with (**A-C**) DMSO, DMSO/Sorbitol (S), Isoxaben (ISX), ISX/S or (**D-F**) boiled Driselase (bDri), bDri/S, Driselase (Dri) and Dri/S. (A, D) Jasmonic acid (JA) and (B, E) Salicylic acid (SA) contents were quantified 7 h after treatment. Values are means (*n* = 4) and error bars represent SD. (C, F) Callose depositions in cotyledons have been quantified 24 h after treatment. Values are means (*n* = 15-20) ± SEM. (**G**) Lignification in Col-0 cotyledons was visualized 24 h after treatment by Phloroglucinol staining. The scale bar represents 1 mm. (**H**) Relative *TOUCH4* (*TCH4)* expression level in Col-0 seedlings 7 h after treatment. Values are means (*n* = 3) ± SD. (**I**) Relative *PLANT DEFENSIN1.2* (*PDF1.2)* expression level in Col-0 and *bak1-5* seedlings 7 h after treatment with bDri or Dri compared to non-treated (NT) seedlings. Values are means (*n* = 3) ± SD. Different letters between treatments of each genotype in (A-I) indicate statistically significant differences according to one-way ANOVA and Tukey’s HSD test (α = 0.05). Asterisks in (A-F) indicate statistically significant differences to the wild type (Student’s *t*-test; ^*^ *P* < 0.05; ^**^ *P* < 0.01; ^***^ *P* < 0.001; ns, not significant).

It is conceivable that a secreted compound, generated by ISX- or driselase-treatment, is responsible for CWD responses. To test this concept, JA/SA levels were measured (i) in *ixr1-1* seedlings, which had been incubated with supernatants from Col-0 seedlings treated with ISX and (ii) in Col-0 seedlings incubated with boiled supernatants deriving from Col-0 seedlings treated with driselase for extended periods (Experimental design summarized in Figure S1B and C). JA and SA levels were barely above the detection limit in the seedlings examined (Figure S1 D, E; compare with Figure 1A-B, D-E). Only minor changes were detected, exemplified by SA levels in seedlings treated with boiled driselase having similar effects as active driselase.

Lignin deposition was detected in Col-0, *bak1-5*, *ixr1-1* cotyledons (red staining, Figure 1G Col-0, Figure S1A *bak1-5* and *ixr1-1*). DMSO, sorbitol and boiled driselase treatments did not cause detectable lignin deposition in the cotyledons examined. In ISX- and driselase-treated Col-0 cotyledons, lignin deposition was detectable in vasculature (ISX) and non-vascular (driselase) tissue (Figure 1G, ISX and Dri). While ISX-treated *ixr1-1* cotyledons did not exhibit increased lignin deposition, the pattern observed in driselase-treated *ixr1* was similar to Col-0 (Figure S1A). Besides *bak1-5* cotyledons exhibiting lignin deposition patterns similar to ISX- or driselase-treated Col-0, both *ixr1-1* and *bak1-5* cotyledons seemed more sensitive to enzymatic digestion by driselase than Col-0 (Figures S1A and 1 G).

Expression levels of *TOUCH4* (*TCH4*) and *PLANT DEFENSIN1.2* (*PDF1.2*) were determined to assess the impact of different CWD types on the expression of reporters for mechanical stimulation (*TCH4*) or defense signaling (*PDF1.2*) (41, 42) (Figure 1H, I). *TCH4* expression was induced in Col-0 by ISX and active driselase while sorbitol co-treatments reduced it, suggesting sensitivity to CWD and/or turgor manipulation (Figure 1H). *PDF1.2* expression was investigated in Col-0 and *bak1-5* seedlings mock-, driselase- or boiled driselase-treated. *PDF1.2* expression was induced similarly in Col-0 seedlings regardless if driselase was active or boiled suggesting, that driselase contains heat-insensitive compounds (possibly PAMPs) inducing *PDF1.2* expression (Figure 1I). *PDF1.2* expression was enhanced in driselase-treated *bak1-5* seedlings but barely induced by boiled driselase, suggesting that two different processes regulate *PDF1.2* expression. One is dependent on BAK1 (boiled driselase sample) while the other one is only detectable / active upon BAK1 loss (active driselase).

Treatments with 300 mM sorbitol normally result in mild hyper-osmotic shocks (43). However, here they reduce the effects of both ISX- and driselase-treatments (Figure 1A-H). This suggests that ISX-/driselase-treatments have opposite effects on turgor levels compared to sorbitol. ISX/driselase seem to generate the equivalent of a hypo-osmotic shock by weakening cell walls while turgor pressure remains high, thus leading to turgor-based cell expansion/CWD. To test this hypothesis we monitored simultaneously epidermal cell shape and viability in seedlings expressing the WAVE131-YFP plasma membrane marker stained with propidium iodide (PI) after mock-, sorbitol-, ISX- or ISX/sorbitol-treatment (44). Here we simplified the experimental setup by using only ISX since in previous experiments ISX and driselase caused qualitatively similar responses and ISX is a better characterized and controllable CWD agent (16). ISX-treatment of seedlings resulted in swelling of epidermal cells and occasional cell death, which were both reduced upon sorbitol co-treatment (Figure S2).

Next, we investigated if genes implicated in mechano- (*MCA1*; *MSCS-LIKE4/5/6/9/10; MSL4/5/6/9/10*), hypo-osmotic *(MSCS-LIKE2/3; MSL2/3, MCA 1*) or hyper-osmotic stress (*ARABIDOPSIS HISTIDINE KINASE1*, *2*/*3*, *4*; *AHK1, AHK2/3, AHK4*) and putatively wound perception (*GLUTAMATE-LIKE RECEPTOR 2.5*; *GLR2.5*) are required for the osmotic suppression observed (45). *GLR2.5* was selected since it exhibits the most pronounced ISX-dependent transcriptional changes off all *GLR* genes analyzed and GLRs have been implicated in wounding response (Figure S4 A, B) (11, 46). We included here also the *theseus1-4* allele (*the1-4*) since it has been recently described as gain of function allele with *the1-4* seedlings exhibiting enhanced responses to CWD similar to previously described THE1 overexpression (*35S::THE1*) seedlings (19) (Merz et al., in press). This allows us to test if stimuli perceived by the THE1 RLK are also affected by osmotic support, enabling us to determine where osmosensitive processes are active in relation to THE1 mediated signaling. Mutant seedlings were mock-, ISX-, sorbitol- or ISX/sorbitol-treated and analyzed for JA accumulation (representative for phytohormone effects) and lignification in the root tip (Figure S3A, B). Only *mca1* and *msl2/3* seedlings exhibited reduced responses to ISX-treatment compared to the corresponding control, while in all genotypes examined osmotic suppression was still detectable. Interestingly ISX/sorbitol-treated *the1-4* seedlings also exhibited reduced responses compared to ISX alone, showing that the stimulus perceived by THE1 is turgor-sensitive. The effects of the osmotic support provided could be based on turgor-equilibration processes (exemplified by the shape changes observed in the epidermal cells, see Figure S2) and would therefore not require any of the sensors tested (47). Turgor manipulation affects all phenotypic CWD effects examined while supernatants from seedlings, which had experienced CWD previously, do not induce phytohormone production. The results of the experiment with *the1-4* seedlings suggest that, whatever stimulus is perceived by the mechanism THE1 is involved in, is also turgor sensitive. Taken together the similarities in seedling responses to different types of CWD (enzymes vs. ISX) suggest that even if the causes of CWD differ, the stimulus activating them may be the same. The results suggest that turgor-sensitive, non-secreted stimuli activate CWD responses. The stimuli could consist of cell wall-bound epitopes changing conformation and/or mechanical distortion/displacement of the plasma membrane against the cell wall upon CWD, similar to the processes activating the CWI maintenance mechanism in *Saccharomyces cerevisiae* (48).

Previously, RLKs required for cell elongation, fertilization and immunity have been implicated in CWI maintenance (2, 8, 17). To gain further insight into the molecular mode of action of the CWI maintenance and to establish, which (if any) of the candidate genes are required, we investigated KO or gain-of-function alleles for 15 RLKs (*THE1*, *CURVY1* (*CVY1*), *FERONIA* (*FER;* knock-down allele *fer-5*), *HERCULES RECEPTOR KINASE1* (*HERK1*), *HERK2*, *ERULUS* (*ERU*), *WAK2/WAK2^cTAP^*, *FEI1*, *FEI2, MIK2*, *BAK1*, *BAK1-LIKE 1* (*BKK1*), *PEPR1*, *PEPR2* and *BIK1)* as well as the *RECEPTOR-LIKE PROTEIN44* (*RLP44*). JA/SA levels were measured in mock- and ISX-treated seedlings of these genotypes and the osmo-/mechano-sensor mutants described above (Figure 2A, B, Figure S5A, B). While mock levels were similar to the corresponding wildtype controls, *fer-5* seedlings exhibited elevated JA/SA levels already in the mock-treated samples, which is in line with the multifaceted functions shown for FER (Figure S5 A-D) (8, 49). Root growth and ISX-resistance were investigated in all lines to exclude pleiotropic effects affecting the analyses performed here (Figure S5E, F). In parallel, we quantified lignin deposition in the root tip area using an image analysis-based approach since for the follow-up data analysis quantitative data was required (Figure 2C). This data for JA, SA, lignin and ISX-resistance (root growth inhibition, RGI) were integrated through phenotypic clustering (Figure 2D). Data for *fer-5* were not included in the clustering to avoid distortion during data integration due to the increased mock hormone levels and because ISX-dependent responses were further increased in *fer-5*, suggesting that *FER* is not essentially required for perception of CWD triggered by ISX (Figure 2A-C, Figure S5C-D). The remaining data (incl. data for osmo-/mechano-sensor mutants, Figure S3) were integrated through phenotypic clustering to generate a global, standardized overview allowing assessment of both relative importance and functions of individual candidates in CWI maintenance (Figure 2D). The results show that KOs in 5 key elements (*BAK1*, *BKK1*, *BIK1*, *PEPR1*, *PEPR2*) PTI cause enhanced responses to ISX-treatment. While the *WAK2^cTAP^* dominant active allele exhibits also slightly elevated responses, *wak2* (Figure S4c), *fei2* and *mik2* seedlings exhibit reductions in the CWD responses examined. KOs in the *Catharantus roseus* RLK1 L (*Cr*RLK1 L) family members *CVY1*, *HERK1* and *HERK2* exhibit enhanced responses whereas *eru* and *the1-1* seedlings exhibit reduced responses, implying functional divergence within the family. Interestingly, loss of *RLP44*, which is involved in cell wall-mediated activation of brassinosteroid signaling, has only very limited effects on the responses suggesting that it is not required for activation of CWD responses (50). In summary, phenotypic clustering shows that WAK2, MIK2, MCA1, MSL2/3, ERU, FEI2 and THE1 are most important for activating CWD responses. Intriguingly they have been previously implicated in turgor/mechano perception and are located in the plasma membrane, plastid envelope or tonoplast, ie. subcellular compartments particularly sensitive to turgor level changes / mechanical stimuli (17, 45). We performed a genetic analysis to establish if THE1, MCA1 and FEI2, which exhibit the most pronounced reductions in CWD responses according to the clustering, are part of the same or different signaling cascades. JA/SA/lignin levels in ISX-treated seedlings were quantified as before (Figure 2E-J). The levels in *mca1 fei2*, *the1-1 mca1* and *the1-1 fei2* seedlings were not additive, but similar to single mutants (Figures 2E-G). *the1-4 mca1* and *the1-4 fei2* seedlings exhibited pronounced reductions in JA/SA levels compared to *the1-4* while relative lignification was only reduced in *the1-4 mca1* (Figure 2H-J). These results suggest that MCA1 and FEI2 are both required for hormone signaling downstream from THE1, but THE1-induced lignification requires only on MCA1.

**Fig. 2.**
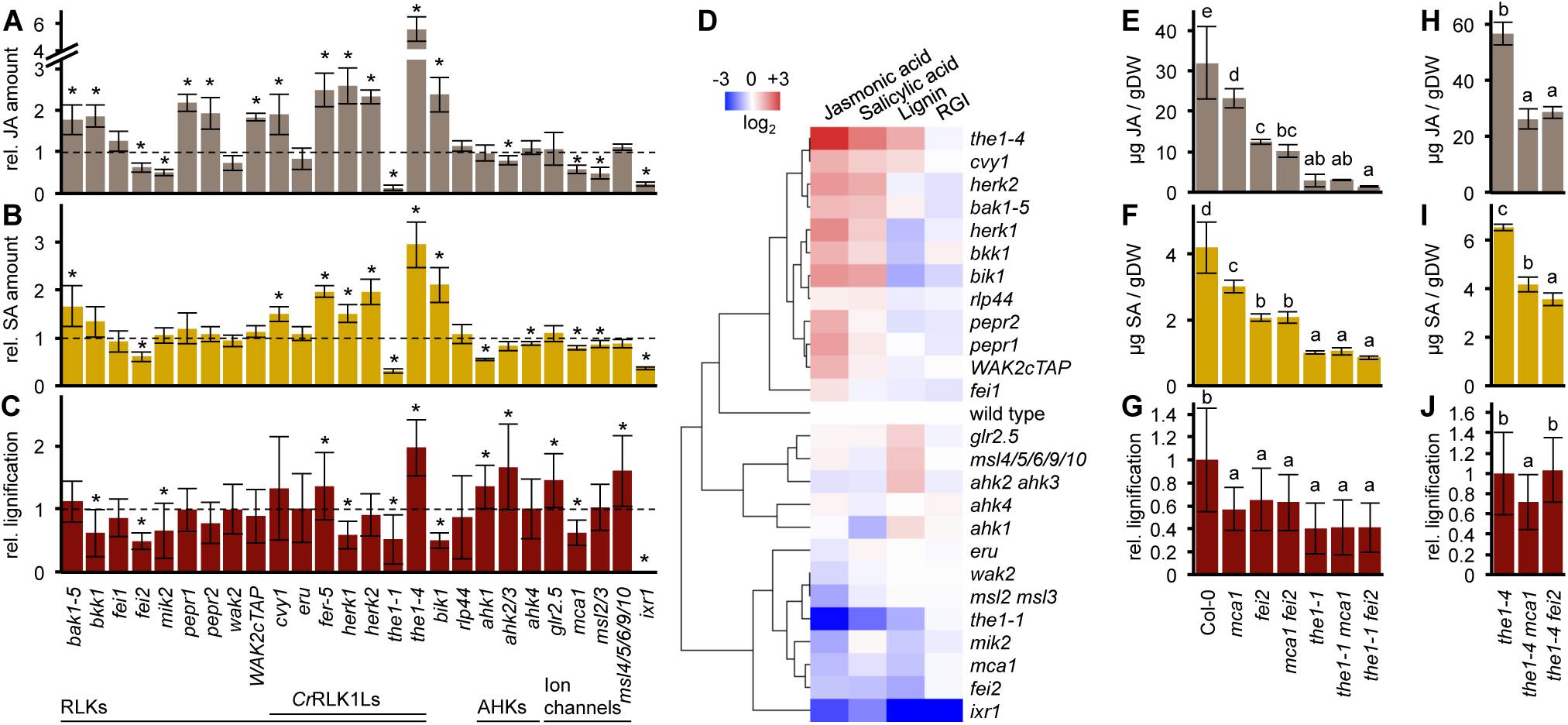
Phenotypic clustering identifies groups of genes involved in cell wall damage responses. (**A**) Jasmonic acid (JA), (**B**) Salicylic acid (SA) and (**C**) root tip lignification were quantified in 6 d-old mutant seedlings after 7 h (A-B; *n* = 4, means ± SD) or 12 h (C; *n* ≥ 10, means ± SD) of Isoxaben (ISX) treatment. Values are relative to a wild type control from a representative experiment selected from at least 3 independent experimental repeats per genotype. Asterisks indicate statistically significant differences to the wild type (Student’s *t*-test, ^*^ *P* < 0.05). Mutant lines are organized in functional groups (RLKs, Receptor-like kinases; *Cr*RLK1 Ls, *Catharanthus roseus* RLK1-like kinases; AHKs, Arabidopsis histidine kinases; Ion channels) and individual genotypes described in detail in Supplemental Table S2. (**D**) Hierarchical clustering of mutant phenotypes assigns functions in CWI maintenance to candidate genes based on their responses to standardized cell wall damage. Mutant phenotype data from (A-C) and Supplemental Figure S5 (RGI, root growth inhibition) have been normalized to wild type controls and log_2_ transformed prior to average linkage clustering. Blue color indicates reduced ISX responses while red color is indicative of increased ISX responses compared to wild type. (**E-G**) Col-0, *mca1, fei2, mca1 fei2, the1-1, the1-1 mca1, the1-1 fei2* and (**H-J**) *the1-4, the1-4 mca1, the1-4 fei2* seedlings were grown for 6 d before treatment with ISX. (E, H) Jasmonic acid (JA) and (F, I) Salicylic acid (SA) were quantified 7 h after treatment (*n* = 4, means ± SD), (G, J) Root tip lignification was quantified 12 h after treatment (*n* ≥17, means ± SD). Different letters between genotypes indicate statistically significant differences according to one-way ANOVA and Tukey’s HSD test (α = 0.05).

To identify novel transcriptionally regulated elements of the CWI maintenance mechanism, Col-0 seedlings were mock- or ISX-treated and analyzed by RNA-Seq (mock- / ISX-treated *ixr1-1* seedlings acted as control). In Col-0 seedlings treated with ISX for 1 hour, 109 transcripts exhibited statistically significant differences from mock-treated controls (p < 0.01, Student’s *t*-test, File S1). None of them was differentially expressed in ISX-treated *ixr1-1* seedlings. GO enrichment analysis detected an overrepresentation of genes implicated in phytohormone-dependent stress responses (Table S1). Amongst the differentially expressed transcripts were also *PROPEP1-4*, which are the precursors for the signaling peptides *At*Pep1-4 (Figure S6) (34). PEPR1 has been shown to bind *At*Pep1-4 while PEPR2 binds only *At*Pep1 and 2 (34). This observation was intriguing since *pepr1* and *2* seedlings exhibit enhanced JA production upon ISX-treatment (Figure 2A). qRT-PCR-based gene expression analysis showed that *PROPEP1* and *3* are particularly strongly induced by ISX (Figure 3A). Time course expression analysis of *PROPEP1* and *3* detected increases in expression levels over time, suggesting that *At*Pep1 and 3 accumulate during the period investigated (Figure 3B). Expression of *PROPEP1* and *3* is still increased in ISX-treated *the1-1* seedlings, suggesting that induction is independent of THE1 mediated processes (Figure 3C). Time course expression analysis of *PROPEP1* and *3* detected increases in expression levels over time, suggesting that *At*Pep1 and 3 accumulate during the period investigated (Figure 3C). To investigate if *At*Pep1 can enhance CWD responses (as described before for JA/SA/ethylene production during wounding) Col-0 seedlings were treated with different concentrations of *At*Pep1 alone or in combination with ISX and JA/SA/lignin production were investigated (Figure 3D-F) (51). *At*Pep1 treatments alone did not affect SA/JA levels, but induced lignin production in a distinctly different pattern than ISX. Surprisingly, seedlings co-treated with ISX/*At*Pep1 exhibited *At*Pep1 concentration-dependent reductions in JA/SA levels, whereas lignin deposition seemed to be additive in ISX/*At*Pep1-treated root tips compared to roots treated with either *At*Pep1 or ISX. To exclude indirect effects and determine if the observed *At*Pep1 effects are mediated via the established *At*Pep1 receptors, the experiments were repeated with Col-0, *pepr1*, *pepr2* and *pepr1/2* seedlings. ISX-induced JA/SA levels were reduced in *pepr1* and *2* seedlings upon co-treatments with *At*Pep1 as observed before (Figure 3G, H). However, this was not the case in *pepr1/2* seedlings, suggesting redundancy and that *At*Pep1 can inhibit ISX-induced phytohormone production via PEPR1 and 2. Interestingly analysis of lignin deposition in *At*Pep1- and/or ISX-treated seedlings showed that PEPR2 (not PEPR1) is essential for *At*Pep1-induced lignin deposition, suggesting differences in signaling activities between PEPR1 and 2 (Figure 3I). These results suggest that CWD activates both the CWI maintenance mechanism but also induces production of *At*PEP1. This production seems to be regulated on the transcriptional level through controlled expression of *PROPEP1* and is independent of THE1-mediated CWD detection. The *At*Pep1 signaling process seems to be redundantly organized because only *pepr1 pepr2* seedlings are unresponsive to *At*Pep1-treatment, whereas the single mutants still do exhibit reductions in ISX-induced phytohormone levels upon ISX/*At*Pep1-cotreatments. Importantly *At*Pep1 seems to act here as an inhibitor of phytohormone accumulation while previously it has been described exclusively as an enhancer of PTI responses. These results suggest that the specific activity of *At*Pep1 is context-dependent.

**Fig. 3.**
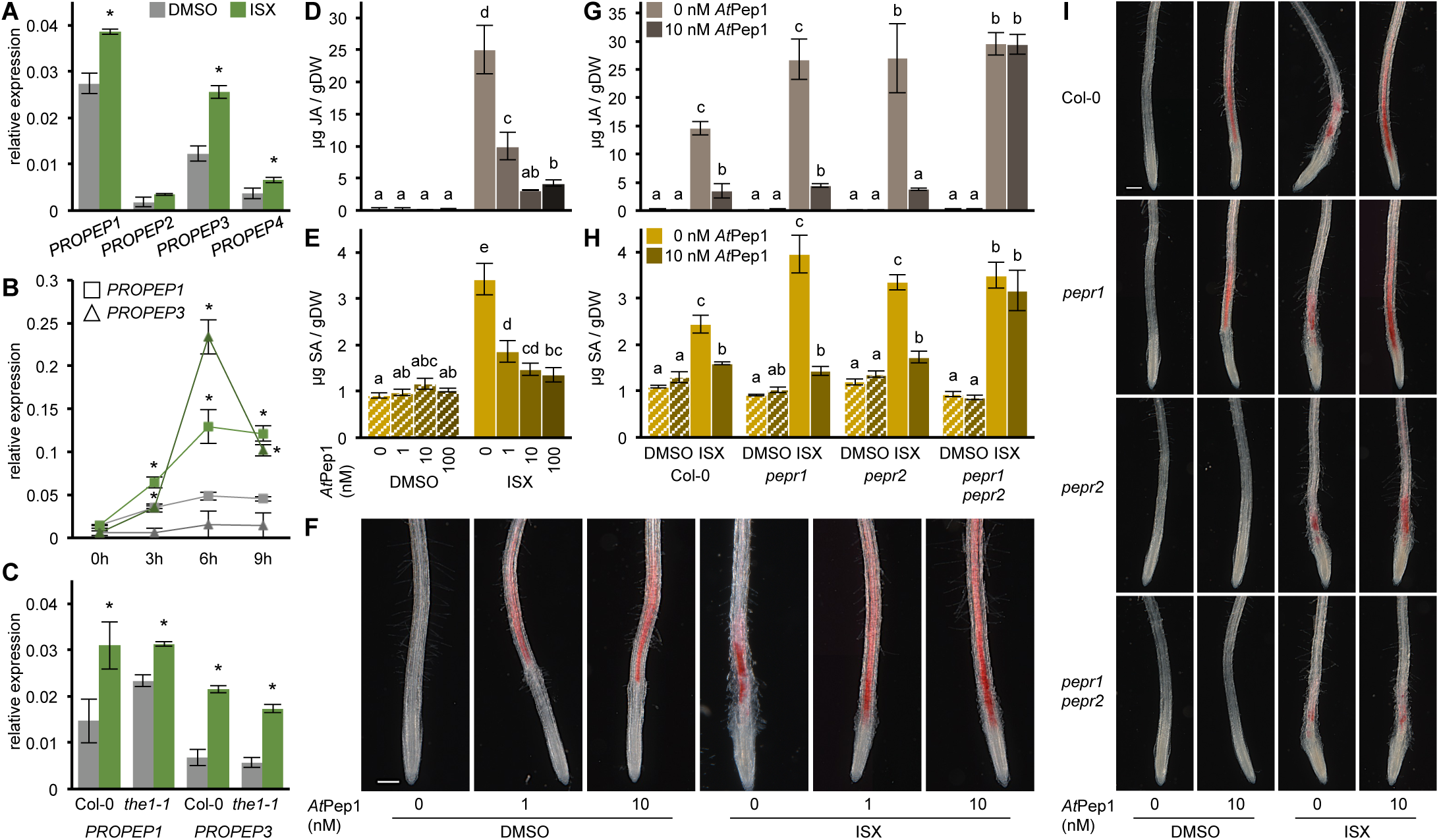
*PROPEP* gene expression is induced by Isoxaben and cell wall damage responses are modulated by co-treatment with *At*Pep1. (**A**) Relative expression levels of *PROPEP1, 2, 3* and *4* were determined by qRT-PCR after 1 h of treatment with DMSO or Isoxaben (ISX) in Col-0 seedlings. *PROPEP1* and *3* expression levels were further examined (**B**) after 0, 3, 6 and 9 h of treatment in Col-0 seedlings and (**C**) after 1 h of treatment with DMSO or ISX in Col-0 and *the1-1* seedlings. Values in (A-C) are means (*n* = 3) ± SD; asterisks indicate statistically significant differences to DMSO-treated controls (Student’s *t*-test; ^*^ *P* < 0.05). (**D**) Jasmonic acid (JA) and (**E**) Salicylic acid (SA) were quantified in Col-0 seedlings after 7 h of co-treatment with DMSO or ISX and 0, 1, 10 or 100 nM *At*Pep1 (*n* = 4, means ± SD). (**F**) Root tip lignification in Col-0 after 12 h of co-treatment with DMSO or ISX and *At*Pep1 (0, 1, 10 nM) was visualized by Phloroglucinol staining. The scale bar represents 200 μm. (**G**) JA and (**H**) SA quantification in Col-0, *pepr1*, *pepr2* and *pepr1 pepr2* after 7 h of co-treatment with DMSO or ISX and 10 nM *At*Pep1 (*n* = 4, means ± SD). Different letters between treatments of each genotype in (D-H) indicate statistically significant differences according to one-way ANOVA and Tukey’s HSD test (α = 0.05). (**I**) Root tip lignification in Col-0, *pepr1*, *pepr2* and *pepr1 pepr2* seedlings after 12 h of co-treatment with DMSO or ISX and 10 nM *At*Pep1 was visualized by Phloroglucinol staining. The scale bar represents 200 μm.

Figure 4 presents a model summarizing how the CWI maintenance mechanism and PTI based processes may regulate early defense responses. Plants apparently perceive CWD separately through the CWI maintenance mechanism and PTI, since *PROPEP1* expression is still induced in *the1-1* seedlings upon treatment with ISX. The CWI maintenance mechanism involves a core group of RLKs and ion-channels located at the plasmamembrane, in the plastid envelope and tonoplast, enabling plant cells to detect mechanical damage to their cell walls or the consequences thereof (Fig. 4, blue elements) (8, 17, 45, 52). The phenotypic data presented here in combination with the molecular functions of the candidate genes identified suggest that the initial stimulus perceived is non-secreted and could be connected to the cell wall or a membrane. The members of this core group probably activate Ca^2+^-based signaling processes, which induce production of ROS, JA and SA (20). Changes in phytohormone and ROS levels in turn modulate downstream responses (exemplified by lignin and callose production). While CWI maintenance has originally been only associated with developmental processes, more recently cell wall signaling and CWI maintenance components have also been implicated in pathogen response in Arabidopsis as well as rice and maize, suggesting the mechanism is active whenever plant CWI is impaired (3, 4, 22). PTI is probably activated by release of *At*Pep1 in response to CWD. This event enhances immune signaling and controls the extent of responses activated by the CWI maintenance mechanism over time (Figure 4, red elements). This control function is supported by results from experiments with *At*Pep1 in ISX-treated seedlings (reduction in CWD responses) and the phenotypes observed with *pepr1*, *pepr2*, *bak1-5*, *bik1* and *bkk1* seedlings (enhanced CWD responses). The experimental data presented here suggest that during early exposure to CWD both mechanisms (PTI and CWI maintenance) are activated in plants. If damage persists (i.e. the CWD is not short term but longer) activation of PTI is enhanced through an *At*Pep1-based positive feedback loop leading to activation of the full immune response (34, 53). Simultaneously the same feedback loop represses the activity of the CWI maintenance mechanism, because the responses are mediated by the full immune response in a specialized manner. If *At*Pep1-mediated PTI is impaired, suppression of CWD-induced responses does not occur, so stress responses continue to be activated by the CWI mechanism. Thus CWI can compensate for defective PTI by causing enhanced phytohormone, lignin, callose accumulation.

**Fig. 4.**
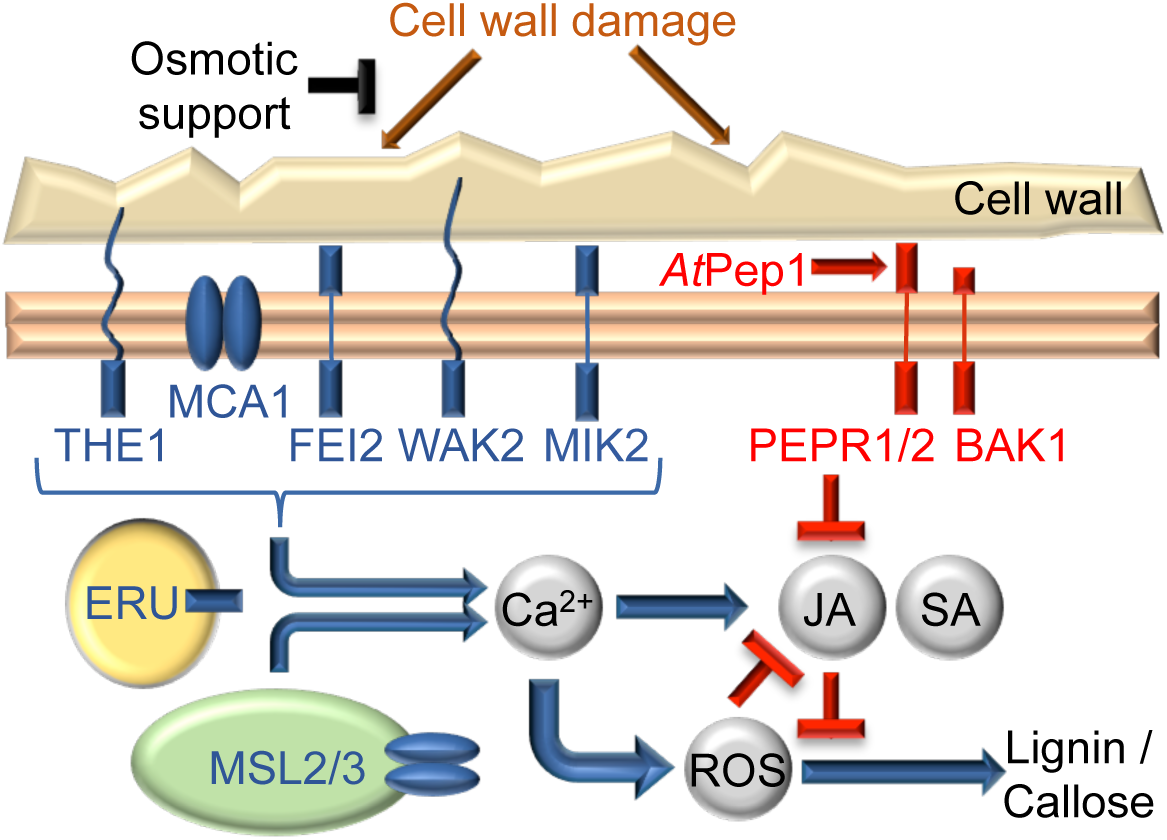
Model of the plant cell wall integrity maintenance mechanism. Responses to cell wall damage in Arabidopsis depend on the receptor-like kinases THE1, FEI2, WAK2, MIK2 and ERU and the ion channels MCA1 and MSL2/3 (Figure 2). THE1, FEI2, WAK2 and MIK2 are localized to the plasma membrane, ERU to the tonoplast and MSL2/3 to plastids (17, 19, 22, 52, 54). Signaling downstream of RLKs and ion channels involves Ca^2+^, reactive oxygen species (ROS) and the phytohormones jasmonic acid (JA) and salicylic acid (SA) (11, 20). Induction of cell wall damage responses can be quantitatively suppressed by provision of osmotic support (11, 55) this study). Cell wall damage also induces the expression of PROPEPs, precursors of elicitor peptides (*At*Pep), independent from THE1-dependent CWD-signaling (Figure 3). RLKs required for Pattern-triggered immunity such as the *At*Pep receptors PEPR1 / 2 and the co-receptor BAK1 repress CWD-induced phytohormone accumulation (Figure 2).

To summarize, the results presented here suggest that PTI and the CWI maintenance mechanism both “detect” CWD in plant cells in different ways and modulate responses in an adaptive manner. Coordination between PTI and CWI maintenance signaling is apparently mediated by *At*Pep1, which functions here as a repressor not an enhancer of downstream responses. The CWI maintenance mechanism seems to act as backup system activating basal broad-spectrum defense in case PTI and activation of the regular defense responses are impaired.

## Author contributions

T.E., N.GB., M.V., T.H. contributed to experimental design, generated data and co-wrote the manuscript. J. McK. and F.A. contributed with experimental data and to the writing of the manuscript. D.D. and C.Z. provided unpublished resources and contributed to the writing of the manuscript.

## Acknowledgements

The authors would like to thank Martin Kuiper and Lauri Vaahtera for constructive comments. Financial and/or technical support from the Deutsche Forschungs-gemeinschaft EN 1071/1-1, HORIZON2020 (SugarOsmoSignaling) (T.E.), BBSRC (J.F. McK.), NTNU (M.V.), EEA grant 7F14155 CYTOWALL (N.G.B./T.H.) and the PROMEC and GCF Facilities at NTNU are gratefully acknowledged. Data generated in the transcriptomics experiments is being deposited with the Bio-analytic Resource for Plant Biology.

---

**Fig. S1. Analysis of hormone accumulation after incubation with supernatants from CWD-treatments and detection of CWD-induced lignification.** (**A, B**) Schemes illustrating the experimental setup to investigate whether compounds secreted upon CWD elicit phytohormone accumulation. Col-0 seedlings have been treated with DMSO, Sorbitol (S), Isoxaben (ISX), Driselase (Dri), boiled Dri (bDri) or combinations thereof for 12 or 24 h. (A) Supernatants from DMSO, ISX and ISX/S treatments have been transferred on 6 d-old *ixr1-1* seedlings. (B) Supernatants from DMSO/S, bDri, bDri/S, Dri and Dri/S have been boiled for 10 min and transferred on 6 d-old Col-0 seedlings.(**C**) Jasmonic acid (JA) and (**D**) Salicylic acid (SA) were quantified 7 h after start of incubation. Values are means (*n* = 4) and error bars represent SD. Different letters between treatments of each genotype and time point indicate statistically significant differences according to one-way ANOVA and Tukey’s HSD test (α = 0.05). (**E**) CWD-induced lignification was analysed in cotyledons of 6 d-old *bak1-5* and *ixr1-1* seedlings. Seedlings were treated with DMSO, Isoxaben (ISX), Driselase (Dri), boiled Dri (bDri) with or without addition of Sorbitol (S). Cotyledons were stained with Phloroglucinol to visualize lignin deposition 24 h after treatment. The scale bar represents 1 mm.

**Fig. S2. CWD-induced root cell bulging is suppressed by sorbitol co-treatment.** 6 d-old WAVE-131YFP seedlings have been treated with DMSO, Sorbitol (S), Isoxaben (ISX) or combinations thereof. After 7 h of treatment images were taken using HC PL APO 10x/0.40 DRY on a Leica SP8. Propidium iodide (PI) staining (1 min, 10 μg/ml) was used to stain the cell walls. The fluorescence related to WAVE-131YFP (green) and PI (red) were used to identify the cell structure. PI and WAVE-131YFP signals largely co-localize in the epidermis and in root hairs (merge). Arrowheads point to areas of ISX-treated roots that are magnified in insets. Scale bar represents 250 μm.

**Fig. S3. ISX responses in candidate Arabidopsis osmo- and mechano-sensor mutant lines are osmo-sensitive.** (**A**) Jasmonic acid (JA) was quantified in 6 d-old *mca1*, *msl2/3*, *msl4/5/6/9/10*, *ahk1*, *ahk2/3, ahk4, glr2.5*, and *the1-4* seedlings after 7 h of treatment with DMSO, Sorbitol (S), Isoxaben (ISX) or combinations thereof. Values are relative to an ISX-treated wild type control (*n* = 3-4, means ± SD). Asterisks indicate statistically significant differences to the wild type (white asterisks) or between treatments (black asterisks; Student’s *t*-test, ^*^ *P* < 0.05, ^**^ *P* < 0.01). (**B**) Col-0, *mca1*, *ahk2/3*, *msl4/5/6/9/10*, *glr2.5*, Ws-2, *ahk1*, *ahk4, msl2/3* and *the1-4* seedlings were stained with Phloroglucinol 12 h after treatment to visualize root tip lignification. Scale bars represent 200 μm.

**Fig. S4: Expression survey of glutamate like receptor (*GLR*) genes and characterization of a *WAK2* T-DNA insertion line.** (**A**) Changes in transcript levels of 18 *GLR* genes in isoxaben-treated Col-0 seedlings after 0, 4, 8, 12, 18, 20, 24, 36 h. Y-axis shows fold change in isoxaben-treated Col-0 seedlings based on normalization to transcript levels in mock-treated seedlings on the Affymetrix ATH-1 chip (Hamann et al., 2009). (**B**) Sketch of the *GLR2.5* gene and results from semi-quantitative RT-PCR reactions for *GLR2.5* compared to *ACT1*. Boxes represent exons, lines introns, a triangle the SALK_016115 T-DNA insertion site and arrowheads primers used for RT-PCR analysis. (**C**) Sketch of the *WAK2* gene and *WAK2* expression relative to *ACT2* determined by qRT-PCR in Col-0 and *wak2-12* seedlings. Boxes represent exons, lines introns, a triangle the SAIL_12_D05 T-DNA insertion site and arrowheads primers used for qRT-PCR analysis. Asterisks indicate a statistically significant difference (Student’s *t*-test; ^***^*P* < 0,001).

**Fig. S5. Analysis of hormone accumulation upon mock-treatment, root length and ISX-resistance in Arabidopsis mutant seedlings.** (**A**) Jasmonic acid (JA) and (**B**) Salicylic acid (SA) were quantified in 6 d-old mutant seedlings after 7 h mock-treatment (DMSO). Columns represent the average of mean values from 3-4 independent experiments, relative to the respective wild type controls (± SD). Mutant lines are organized in functional groups (RLKs, Receptor-like kinases; *Cr*RLK1 Ls, *Catharanthus roseus* RLK1-like kinases; AHKs, Arabidopsis histidine kinases; Ion channels; cf. Fig.2) and individual genotypes described in detail in Supplemental Table S2. Absolute amounts of (**C**) JA and (**D**) SA in Col-0 and *fer-5* seedlings treated with DMSO or ISX for 7 h illustrate increased basal hormone levels in *fer-5*. (**E**) Root length and (**F**) ISX-resistance (root growth inhibition, RGI) assays show growth defects and no resistance to ISX-treatment in different mutant seedlings. To assess growth phenotypes, root lengths of 6 day-old wild type and mutant seedlings were measured prior to treatment. Resistance to ISX was determined 24 h after treatment. 100% indicates full sensitivity to ISX-treatment, while 0% indicates complete resistance. Columns represent the average of mean values from 3-8 independent experiments ± SD. Different letters between genotypes in (A, B, E, F) indicate statistically significant differences according to one-way ANOVA and Tukey’s HSD test (α = 0.05). Asterisks in (C-D) indicate statistically significant differences to the wild type (Student’s *t*-test, ^*^ *P* < 0.05).

**Fig. S6. Changes in *PROPEP* gene expression upon ISX-treatment.** 6 d-old Col-0 seedlings were treated with DMSO or ISX. Changes in gene expression were analyzed after 1 h using RNA-Seq (see Supplemental File S1). ISX-induced changes in gene expression of *PROPEP1, 2, 3, 4, 5, 6* and *7* are plotted on a log_2_ scale and *P*-values are given next to the individual bars (*n* = 3).

**Table S1. Gene Ontology (GO) enrichment analysis of genes with ISX-dependent expression changes.** Genes with significantly altered expression after 1 h of ISX-treatment (Supplemental File S1) have been analyzed for GO enrichment using the PANTHER Overrepresentation Test (release 20160715) and the GO Ontology database (Released 2017-02-28) on http://geneontology.org/. Results are filtered by *P* < 0.05 after Bonferroni correction for multiple testing.

**Table S2. Arabidopsis genotypes used in this study.** Names used in this work are listed along with gene identifiers, specific allele used, wild type background and references.

**Table S3. Primers used in this study.**

**File S1. RNA-Seq analysis of gene expression changes upon 1 h of ISX-treatment.** 6-day old Col-0 and *ixr1-1* seedlings were treated with DMSO or ISX and analyzed by RNA-Seq 1 h after treatment. Raw data from 3 independent biological replicates are shown in the “raw data” tab. 109 transcripts found differentially regulated after ISX-compared to DMSO-treatment (p< 0.01) are shown in the tab “Col ISX vs DMSO”. Up- or downregulated transcripts compared to DMSO controls are color-coded orange (up) or blue (down). Fold changes of *PROPEP1, 2, 3, 4, 5, 6* and *7* transcripts are calculated in tab “PROPEP ISX vs DMSO” and plotted in Supplemental Fig. S6.

## Materials and Methods

### Plant growth and treatment

*Arabidopsis thaliana* genotypes used in this study were obtained from the labs previously publishing them, or ordered from the Nottingham Arabidopsis Stock Centre (http://arabidopsis.info/). Detailed information is listed in Supplemental Table S2. Seedlings were grown in liquid culture as described by (20) with minor modifications. 30 mg of seeds were sterilized by sequential incubation with 70 % ethanol and 50 % bleach on a rotating mixer for 10 min each and washed 3 times with sterile water. Seeds were than transferred into 250 ml erlenmayer flasks containing 125 ml half-strength Murashige and Skoog growth medium (2.1 g/L Murashige and Skoog Basal Medium, 0.5 g/L MES salt and 1 % sucrose at pH 5.7). Seedlings were grown in long-day conditions (16 h light, 22°C / 8 h dark, 18°C) at 150 μmol m^-2^ s^-1^ photon flux density on a IKA KS501 flask shaker at a constant speed of 130 rotations per minute. Seedlings were treated after 6 days with 600 nM isoxaben (ISX; in DMSO), 0.03% (w/v) driselase (Dri), 0.03% Dri boiled for 10 min, 300 mM sorbitol (S) and combinations thereof in fresh growth medium. Supernatants from treated Col-0 cultures were incubated with *ixr1-1* seedlings (DMSO, ISX, ISX/S) or boiled for 10 min and incubated with Col-0 seedlings (DMSO/S, bDri, bDri/S, Dri, Dri/S). *At*Pep1 peptide (ATKVKAKQRGKEKVSSGRPGQHN) was obtained from Peptron (Daejeon, South Korea) and dissolved in sterile water.

### Phytohormone Analysis

Jasmonic acid (JA) and salicylic acid (SA) were analyzed as described by (Forcat et al. 2008) with minor modifications. Seedlings were flash-frozen in liquid nitrogen and freeze-dried for 24 h. 6-7 mg aliquots of freeze-dried seedlings were ground with 5 mm stainless steal beads in a Qiagen Tissue Lyser II for 2 min at 25 Hz. Shaking was repeated after addition of 400 μl extraction buffer (10 % methanol, 1 % acetic acid) with internal standards (10 ng Jasmonic-d_5_ Acid, 28 ng Salicylic-d_4_ Acid; CDN Isotopes, Pointe-Claire, Canada) before samples were incubated on ice for 30 min and centrifuged for 10 min at 16,000 g and 4°C. Supernatants were transferred into fresh tubes and pellets re-extracted with 400 μl extraction buffer without internal standards. Supernatants were combined and centrifuged 3 times to remove all debris prior to LCMS/MS analysis. An extraction control not containing plant material was treated equally to the plant samples.

Chromatographic separation was carried out on a Shimadzu UFLC XR, equipped with a Waters Cortecs C18 column (2.7 μm, 2.1 × 100 mm). The solvent gradient (acetonitrile (ACN) / water with 0.1 % formic acid each) was adapted to a total run time of 7 min: 0-4 min 20 % to 95 % ACN, 4-5 min 95 % ACN, 5-7 min 95 % to 20 % ACN; flow rate 0.4 ml / min. For hormone identification and quantification an AB SCIEX Triple Quad 5500 system was used. Mass transitions were: JA 209 > 59, D_5_-JA 214 > 62, SA 137 > 93, D_4_-SA 141 > 97.

### Callose Analysis

Seedlings were sampled 24 h after treatment and placed in 70 % (v/v) ethanol. For callose staining, samples were incubated in 0.07 M sodium phosphate buffer pH 9 for 30 min and in 0.005 % (w/v) aniline blue (in 0.07 M sodium phosphate buffer pH 9) for 60 min. Samples were washed with water, mounted in 50 % (v/v) glycerol and analyzed under a Nikon Eclipse E800 microscope using a UV-2A filter (EX 330-380 nm, DM 400 nm, BA 420 nm). Images were taken at 10x magnification and callose depositions quantified using ImageJ software.

### Lignin Analysis

Lignification was investigated 12 h (root tips) and 24 h (cotyledons) after start of treatments. Lignin was detected with phloroglucinol-HCL as described (Denness et al., 2011). Seedlings were photographed using a Zeiss Axio Zoom.V16 stereomicroscope. To assess the extent of lignin production in root tips, phloroglucinol-stained areas and the total root area imaged were quantified using ImageJ (the same root length was maintained in all images taken). The relative lignified area was plotted as fold change compared to wild type root tips.

### Root Measurements

Absolute root lengths were measured immediately prior to ISX treatment (0 h) to examine root growth phenotypes and 24 h after start of treatment to determine ISX-dependent root growth inhibition (RGI). For calculation of %RGI the following formula was applied: [1 - (ISX 24h - ISX 0h) / (mock 24h - mock 0h)]^*^100.

### Hierarchical Cluster Analysis

Hierarchical Clustering of ISX-dependent phenotypes was performed with Cluster 3.0 using the C Clustering Library v1.52 (de Hoon et al., 2004). All data of mutant seedlings was normalized to their corresponding wild type control. Log_2_ transformed data was then used for average linkage clustering with an uncentered correlation similarity metric. Results were depicted using Java TreeView v1.1.6r4 (Saldanha 2004) and color-coded blue (less than in wild type) or red (more than in wild type).

### Semi-quantitative RT-PCR

Total RNA was isolated using a Qiagen RNeasy Mini Kit in combination with DNaseI treatment according to manufacturer instructions (www.quiagen.com). For reverse Sequences of primers used for amplification (*ACT1, GLR2.5*) are listed in Supplemental Table S3.

### qRT-PCR

Total RNA was isolated using a Spectrum Plant Total RNA Kit (Sigma-Aldrich). 2 μg of total RNA were treated with RQ1 RNase-Free DNase (Promega) and processed with the ImProm-II Reverse Transcription System (Promega) for cDNA synthesis. qRT-PCR was performed using a LightCycler 480 SYBR Green I Master (Roche) and primers (Supplemental Table S3) diluted according to manufacturer specifications. Four different reference genes (*PP2A, ACT2, UBA1, GRF2*) were examined to identify one exhibiting stable expression during ISX-treatment. *ACT2* was the most stable one and used in all experiments as reference.

### RNA-Seq

Total RNA was extracted using a Spectrum Plant Total RNA Kit (Sigma-Aldrich). RNA concentration was measured using a Qubit RNA HS Assay Kit (Thermo Fisher Scientific) and integrity assessed using a Agilent RNA 6000 Pico Kit. RNA Seq libraries were prepared using a TruSeq Stranded mRNA Kit (Illumina) according to the manufacturer’s instructions. 500 ng total RNA was used as starting material. First, index barcodes were ligated for identification of individual samples. mRNA purification, fragmentation and cDNA synthesis has been performed as described in (Ren et al. 2015). Exonuclease / polymerase was used to produce blunted overhangs. Illumina SR adapter oligonucleotides were ligated to the cDNA after 3’ end adenylation. DNA fragments were enriched by 15 cycles of PCR reaction. The libraries were purified using the AMPure XP (Beckman Coulter), quantitated by qPCR using a KAPA Library Quantification Kit (Kapa Biosystems) and validated using a Agilent High Sensitivity DNA Kit on a Bioanalyzer. The size range of the DNA fragments were measured to be in the range of 200-700 bp and peaked around 296 bp. Libraries were normalized and pooled to 2.2 pM and subjected to clustering on NextSeq 500 high output flowcells. Finally single read sequencing was performed for 75 cycles on a NextSeq 500 instrument (Illumina) according to the manufacturer instructions. Base-calling has been performed on the NS500 instrument by Illumina RTA v2.4.6. FASTQ files were generated using bcl2fastq2 Conversion Software v1.8.4. Each FASTQ file was subjected to quality control trough fastQC v11.1 before technical replicates were combined and an average of 13.1 million reads was produced for each library. The reads were then aligned to the *A. thaliana* genome (Ensembl v82) with STAR v2.4.1 in two-pass mode. On average, 96.2% of the reads aligned to the genome. The reads that aligned uniquely to the genome were aggregated into gene counts with FeatureCounts v1.4.6 using the genome annotations defined in Ensembl v82. Of the 32000 genes defined in the gene model, a total of 20750 genes were left for analysis after filtering out genes with a CPM (counts-per-million) value less than one in two or more samples.

The filtered gene count table was used as input to the Voom method (Law et al. 2014) of the limma R package v3.26.9 for differential expression. The samples were normalized using the TMM (Robinson et al. 2010) method before a linear model was defined. Differential expression between groups were tested by empirical Bayesian moderated *t*-tests and *P*-values were corrected for multiple testing by the Benjamini-Hochberg false discovery rate adjustment. Statistical significance of pairwise comparisons was determined using a Student’s *t*-test.

### Statistical Analysis

Statistical significance was assed using either Student’s *t*-test or one-way ANOVA followed by post-hoc analysis with Tukey’s HSD test. Statistical details of experiments are specified in the figure legends. Statistically significant differences are indicated by ^*^ *P* < 0.05, ^**^ *P* < 0.01, ^***^ *P* < 0.001 for Student’s *t*-test and different letters for one-way ANOVA / Tukey’s HSD test at α = 0.05. All statistical analyses were performed in IBM SPSS Statistics v24.

